# Recombination-aware Phylogeographic Inference Using the Structured Coalescent with Ancestral Recombination

**DOI:** 10.1101/2022.02.08.479599

**Authors:** Fangfang Guo, Ignazio Carbone, David A. Rasmussen

**Affiliations:** Department of Entomology and Plant Pathology, North Carolina State University, Raleigh, North Carolina, USA; Center for Integrated Fungal Research, North Carolina State University, Raleigh, North Carolina, USA; Bioinformatics Research Center, North Carolina State University, Raleigh, North Carolina, USA

**Keywords:** Phylogeography, Migration, Structured coalescent, Recombination, Ancestral recombination graph, *Aspergillus flavus*

## Abstract

Movement of individuals between populations or demes is often restricted, especially between geographically isolated populations. The structured coalescent provides an elegant theoretical framework for describing how movement between populations shapes the genealogical history of sampled individuals and thereby structures genetic variation within and between populations. However, in the presence of recombination an individual may inherit different regions of their genome from different parents, resulting in a mosaic of genealogical histories across the genome, which can be represented by an Ancestral Recombination Graph (ARG). In this case, different genomic regions may have different ancestral histories and so different histories of movement between populations. Recombination therefore poses an additional challenge to phylogeographic methods that aim to reconstruct the movement of individuals from genealogies, although also a potential benefit in that different loci may contain additional information about movement. Here, we introduce the Structured Coalescent with Ancestral Recombination (SCAR) model, which builds on recent approximations to the structured coalescent by incorporating recombination into the ancestry of sampled individuals. The SCAR model allows us to infer how the migration history of sampled individuals varies across the genome from ARGs, and improves estimation of key population genetic parameters such as population sizes, recombination rates and migration rates. Using the SCAR model, we explore the potential and limitations of phylogeographic inference using full ARGs. We then apply the SCAR to lineages of the recombining fungus *Aspergillus flavus* sampled across the United States to explore patterns of recombination and migration across the genome.

## Introduction

In the absence of any recombination, populations evolve clonally and the ancestral relationships among all individuals can be captured by a single genealogy or phylogenetic tree [Rosenberg and Nordborg, 2002, Hein et al., 2005]. However, in the presence of recombination, individuals can inherit different parts of their genome from different ancestors, leading to a mosaic of phylogenetic relationships across the genome that cannot be captured by any single tree. Since many population genetic and phylogeographic methods infer demographic parameters (e.g. population sizes, migration rates) from a phylogeny assumed to reflect the clonal ancestry of sampled individuals, recombination poses a major challenge to demographic inference.

Rates of recombination vary dramatically from asexual populations that experience no recombination to sexually outcrossing populations where recombination occurs between parental genomes every generation [Smith et al., 1993, Stapley et al., 2017, Hasan and Ness, 2020]. How demographic inference methods deal with recombination largely depends on the assumed rate of recombination. If recombination rates are very low, individuals will inherit large regions of their genome (i.e. non-recombinant blocks) from the same set of ancestors. Moreover, recombination will only impact the ancestry of lineages directly involved in a recombination event while preserving the ancestral relationships among non-recombinant lineages [Hein et al., 2005]. In this case, phylogenies can be reconstructed from non-recombinant regions of the genome or recombining lineages can be identified and removed. At the other extreme, very high rates of recombination will break apart linkage between loci, such that different loci can be treated independently [Hudson, 1990]. In this case population genomic methods that treat each locus as (pseudo-)independent can be used to draw demographic inferences [Speidel et al., 2019].

However, in between these two extremes lie many organisms that undergo intermediate amounts of recombination, including many important microbial pathogens [Goss, 2015]. For example, this includes many fungi with mixed mating systems that are predominately clonal but occasionally reproduce sexually and thus recombine [Nieuwenhuis and James, 2016]. Such intermediate rates of recombination pose a particular challenge to demographic inference as it may be difficult to identify and accurately reconstruct phylogenetic relationships from any non-recombinant genomic region. At the same time, recombination is not frequent enough to breakdown correlations among linked loci, violating assumptions of independence between loci and simply concatenating alignments may lead to phylogenetic reconstructions inconsistent with the true ancestry of the sample [Kubatko and Degnan, 2007].

Ideally, the differing but correlated patterns of ancestry across the genome would be captured using ancestral recombination graphs (ARGs) [Griffiths and Marjoram, 1997]. An ARG describes the complete genealogical history of sampled individuals, including *local* trees representing the genealogy of sampled individuals over a particular non-recombination region of the genome and the recombination events connecting lineages across local trees. Although reconstructing full ARGs is notoriously difficult, recent advances now allow ARGs to be accurately reconstructed for a modest number of samples (i.e. *<*100). Notably, ARGweaver allows for full Bayesian inference of ARGs under the sequential Markov coalescent (SMC) model, an approximation to the full coalescent with recombination [Rasmussen et al., 2014b]. More recent methods allow for ARGs to be approximated for much larger datasets by reconstructing series of local or marginal trees using approximate methods [Kelleher et al., 2019, Speidel et al., 2019].

In addition to recombination, population structure can also strongly shape the genealogical history of a population. The structured coalescent extends basic coalescent models by allowing lineages to migrate between different subpopulations or demes [Notohara, 1990]. While the structured coalescent is most often used to model geographic structure, the theory holds for many different forms of population structure (e.g. assortative mating within a population) [Wakeley, 2009]. Under the structured coalescent, migration rates can be estimated from a genealogy of individuals sampled from different populations [Beerli and Felsenstein, 2001], and structured coalescent models form the basis of several phylogeographic inference frameworks [Maio et al., 2015, Müller et al., 2018, Vaughan et al., 2014]. However, population structure also poses a major challenge to demographic inference under coalescent models because lineages in the genealogy are no longer exchangeable in the sense that the probability of two lineages coalescing will depend on their ancestral location or state. Statistical inference under the structured coalescent therefore requires the ancestral state of lineages to be imputed, and early methods implemented algorithms to sample ancestral states using Markov chain Monte Carlo (MCMC) or other sampling-based methods [Beerli and Felsenstein, 2001]. Because jointly estimating the ancestral locations of all lineages along with the demographic parameters of interest poses yet another computational challenge, more recent methods make various approximations to the full structured coalescent to track the movement of lineages probabilistically, such that the unknown ancestral locations can be marginalized or integrated over [Volz, 2012, Rasmussen et al., 2014a, Müller et al., 2017].

Given that the statistical and computational performance of ARG reconstruction methods will likely continue to improve in coming years, we explore phylogeographic inference where the ARG is assumed to be known or at least reconstructed accurately. We first develop a new model we call the Structured Coalescent with Ancestral Recombination (SCAR) to estimate demographic parameters in the presence of both migration and recombination from a reconstructed ARG. The SCAR model combines recent advances in modeling the coalescent with recombination using the SMC approximation with approximations to the structured coalescent that marginalize over unknown ancestral states [Volz, 2012, Rasmussen et al., 2014a, Müller et al., 2017]. Next, we test the limits of reconstructing ARGs from genomic sequence data using ARGweaver and then explore how accurately demographic parameters can be inferred from reconstructed ARGs using the SCAR model. Using simulated sequence data, we show that parameters such as recombination rates, migration rates and population sizes can be accurately estimated from ARGs under the SCAR model as long as the underlying ARG can be accurately reconstructed. We then apply this approach to the plant fungal pathogen *Aspergillus flavus* to estimate recombination and migration rates between natural populations in several US states.

## Models and Methods

### The SCAR model

#### Description of the SCAR model

Here we incorporate recombination and population structure simultaneously into the basic coalescent process. We consider a population divided into *q* different demes or (sub)populations. Each population *k* is composed of *N*_*k*_ haploid individuals which reproduce each generation to generate a random number of offspring (i.e. a Wright-Fisher population). Two lineages can exchange genetic material through recombination, in which case their children may inherit genetic material from both parents. Lastly, we allow individuals to transition or migrate between populations.

Three different types of events can therefore occur in the ancestry of sampled lineages under this model: coalescent events, recombination events and migration events (see Figure 1). We begin by considering the rate at which each one of these events will occur among lineages in the genealogy.

**Figure 1.**
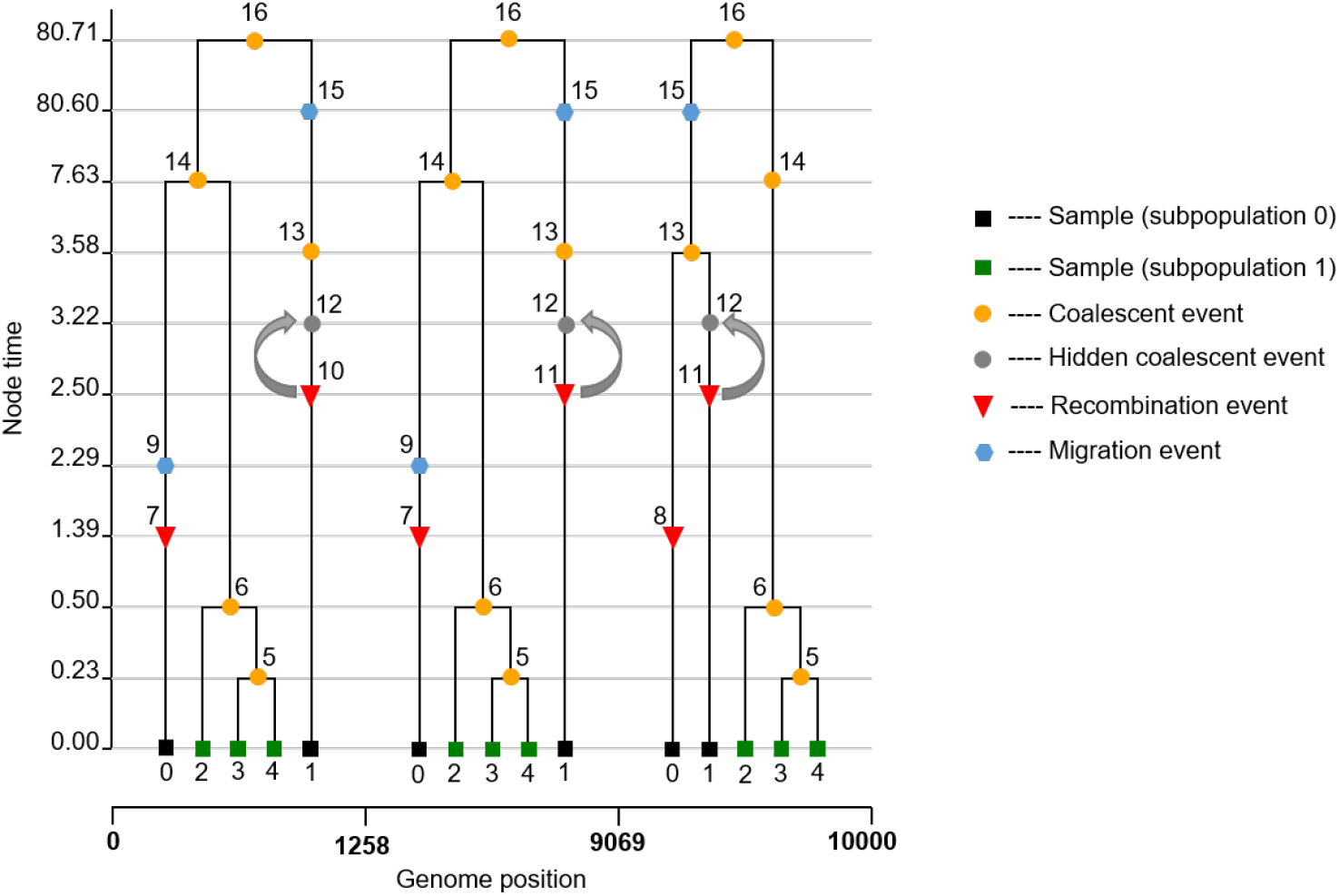
An ancestral recombination graph. In this example, an ARG was simulated for five individuals sampled in two subpopulations (0 and 1) using msprime [Kelleher et al., 2016]. Two recombination events happen, dividing the genome into three segments each with their own local tree. The first and second local tree have the same topology, because after the recombination event (indicated by nodes 10 and 11) the two parent lineages coalesce with one another (indicated by the grey arrow) at a hidden coalescent event (indicated by node 12), which would not normally be observed in the local trees. The second and the third local tree are topologically discordant due to a recombination event (indicated by nodes 7 and 8).

#### Coalescent events

As under the standard coalescent for a Wright-Fisher population [Watterson, 1975, Kingman, 1982], the probability of two lineages finding their most recent common ancestor in a given generation is inversely proportional to the population size. Pairs of lineages in population *k* will therefore coalesce at rate 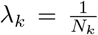 per generation. For now, we will assume lineages in different populations cannot coalesce, although this assumption can be relaxed (see for example Volz [2012]). The total coalescent rate among all pairs of lineages in population *k* is 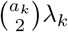, where *a*_*k*_ is the number of lineages in *k*. For now we will also assume the ancestral location of lineages is known.

#### Recombination events

As in earlier models for the coalescent with recombination, each lineage in the tree undergoes a recombination event at rate *r* per site along a genome of length *L* [Hudson, 1990, Kuhner et al., 2000, Kelleher et al., 2016]. However, because two parents contribute genetic material to a child lineage at a recombination event, not all of the genetic material each parent lineage carries will be ancestral to the sample [Hudson, 1990]. Further, because recombination involving non-ancestral genetic material will have no effect on the genealogy of sampled individuals, we only need to consider recombination events involving ancestral genetic material. Following Kuhner et al. [2000], we use the term eligible links to refer to sites that are eligible to undergo recombination because they contribute genetic material ancestral to the sample and occur next to at least one other site eligible to undergo recombination. We therefore track the number of eligible links *B*_*i*_ carried by each lineage *i*, such the total rate at which lineage *i* recombines is *rB*_*i*_. We can then compute the total recombination rate among lineages in population *k* as 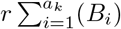.

#### Migration events

Lineages migrate between populations *k* and *l* at rate *γ*_*kl*_ in forwards time. The total rate at which all lineages in population *k* migrate to another population is therefore 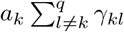.

As in other coalescent models, the time to the next event of each type is exponentially distributed according to the rate of each event type. Furthermore, we assume the coalescent, recombination and migration processes are independent conditional upon the number of lineages *a*_*k*_ in each population state. That is, while events may change the number of lineages in each state, the different events do not influence the probability of the other events occurring over time intervals in which *a*_*k*_ is constant. The three processes are therefore independent, competing processes where the time to the next event is exponentially distributed according to the total rate Ω at which events of any type occur:

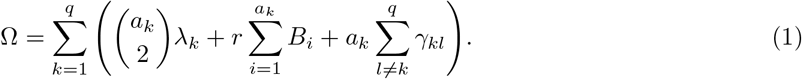

#### The likelihood of an ARG under the structured coalescent with known ancestral states

We now consider how to compute the likelihood of a fully known ancestral recombination graph 𝒢, which can be thought of as a sequence of local trees with additional nodes representing recombination events that split ancestral genetic material among different parent lineages in the local trees. Going backwards in time, at a coalescent event two lineages in state *k* merge into a single parent and the total number of lineages *a*_*k*_ in the ARG in state *k* decreases by one. At a recombination event, a lineage divides into two parent lineages and *a*_*k*_ increases by one. We seek to compute the likelihood *L*(𝒢 | *θ*) of 𝒢 under the SCAR model given a set of demographic parameters *θ* = { *λ*,r, *γ* }, allowing for likelihood-based inference of these parameters from an ARG.

For an ARG with *e*_*c*_ coalescent events, *e*_*r*_ recombination events, and *e*_*m*_ migration events, there will be a total of *e* = *e*_*r*_ + *e*_*c*_ + *e*_*m*_ events in the graph. The ARG can be divided into *e tree intervals*, within which the total number of lineages in the ARG (and in each population) remains the same. We will let 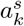 be the number of lineages present in the tree during the *s*-th tree interval. We denote the waiting time between each event as Δ*t*_*s*_ = *t*_*s*_ *− t*_*s−*1_.

Assuming exponentially distributed waiting times between events, the coalescent likelihood has the general form:

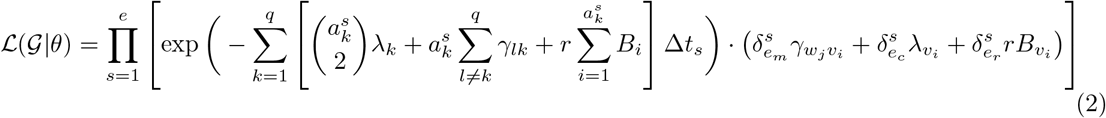

The exponential term gives the probability that in the *s*th time interval with duration Δ*t*_*s*_ no coalescent, recombination or migration event occurs in any population. The remaining term is the point probability density of the event that terminates the interval. We use the indicator variables 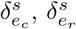 and 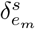 to indicate whether the event terminating interval *s* is a coalescent, migration or recombination event, respectively; where 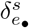 is 1 when the corresponding event type terminates the interval and 0 otherwise.

The events in genealogy 𝒢 are either migrations from subpopulation *w*_*j*_ to *v*_*i*_, or coalescence or recombination events in subpopulation *v*_*i*_.

#### The likelihood of an ARG with unknown ancestral states

Because we typically do not observe the ancestral location or state of lineages, they must either be jointly inferred along with the other model parameters or integrated (marginalized) out when computing the likelihood of the ARG. Here, we use the approximation first proposed by Volz [2012] to track the ancestral state of lineages probabilistically, and then marginalize over ancestral states using these lineage state probabilities.

With unknown ancestral states, the rate at which a pair of lineages *i* and *j* coalesce now depends on the probability that both lineages are in the same population:

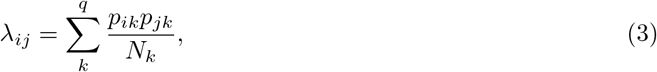

where *p*_*ik*_ and *p*_*jk*_ are the probabilities that lineage *i* and lineage *j* are in state *k*, respectively. How these lineage state probabilities are computed is explained further below.

The total rate at which all lineages *a* coalesce can then be computed by summing over all pairs of lineages:

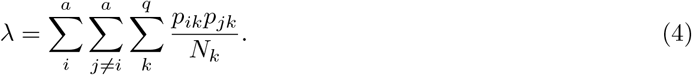

However, repeatedly summing over all possible pairs of lineages can become computationally burdensome, especially as the number of lineages grows large. To avoid this, we can approximate the number of lineages in each state using the lineage state probabilities:

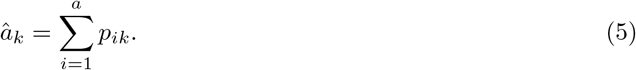

We then approximate the total rate at which pairs of lineages coalesce in state *k* as:

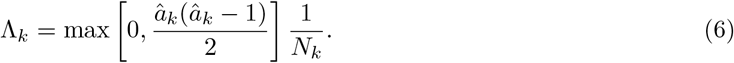

We then compute the total recombination rate in state *k* as:

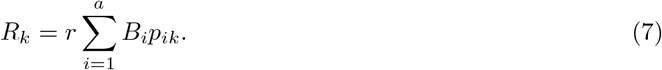

The total likelihood of the ARG when integrating over ancestral states then becomes:

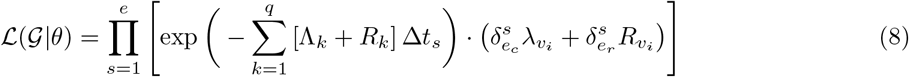

Note that while the migration rates do not directly enter into likelihood function they influence the lineage state probabilities *p*_*ik*_ that in turn determine Λ_*k*_ and *R*_*k*_.

#### Tracking lineage state probabilities

Going backwards in time, a lineage currently residing in population *k* will migrate to population *l* at rate *γ*_*lk*_. Assuming the probability of a lineage residing in a population is independent of the location of all other lineages, the migration process along each lineage can be modeled as a continuous time Markov process on a discrete state space [Volz, 2012]. We can then use a system of differential equations to track how the probability of a lineage residing in each state changes backwards through time:

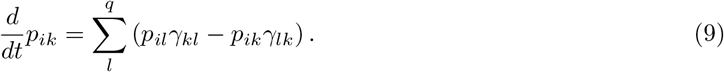

Given a vector of initial lineage state probabilities *p*_*i*_(0) at time zero, we can analytically solve (9) above for *p*_*i*_(*t*) at some time *t* further in the past:

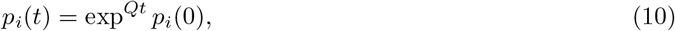

where the matrix *Q* is that transition rate matrix derived from *γ*:

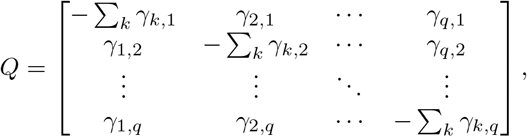

where the forward-time migration rate matrix has been transposed to obtain the backward time rates.

As originally shown in [Müller et al., 2017], these equations are approximate because they assume all lineages evolve independently such that the probability of one lineage residing in a population is completely independent. In contrast, under the exact structured coalescent model, lineages states may be correlated because the observation that two lineages have or have not coalesced can be informative about their location. For example, two or more lineages are unlikely to reside in the same population over long periods of time and not coalesce if *N*_*k*_ is small in population *k*, such that the observation that the lineages have not coalesced increases the probability of these lineages being in different populations. The bias introduced by ignoring the non-independence of lineages is most extreme when: 1) migration rates are low relative to coalescent rates and 2) either coalescent or sampling fractions are highly asymmetric between populations [Müller et al., 2017]. In these cases, a more accurate approximation to the structured coalescent exists but computing lineages state requires solving a high-dimensional system of differential equations. We therefore continue to assume independence among lineages but note that this more complex approximation can be substituted when necessary.

#### Statistical inference under the SCAR model

We use a Bayesian MCMC approach to infer the posterior distribution of demographic parameters. In particular, we use a Metropolis-Hastings algorithm to sample from the joint posterior distribution of parameters given a fixed ARG 𝒢:

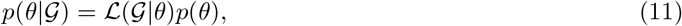

where the likelihood ℒ (𝒢 | *θ*) is computed as in (8) and *p*(*θ*) is the prior distribution on the demographic parameters. In simulation experiments, we chose a uniform distribution for *p*(*θ*) such that our estimates are not influenced by the prior but use informative priors when performing inference from real data.

#### ARG reconstruction using ARGweaver

We use ARGweaver [Rasmussen et al., 2014b] to reconstruct ARGs from sampled genomic sequence data. ARGweaver uses the SMC approximation to compute the likelihood of an ARG evolving under the coalescent with recombination. The coalescent likelihood of the ARG is then combined with the likelihood of the sequence data evolving along each local tree in the ARG to compute the joint likelihood of the sequence data and ARG. ARGweaver then employs a Bayesian MCMC approach to sample ARGs from the corresponding posterior distribution. To obtain a single, representative ARG, we choose the ARG from the posterior sample with either the maximum joint likelihood, maximum (sequence) likelihood, or from the final MCMC iteration.

ARGs sampled by ARGweaver are output in either SMC or ARG format. To facilitate computing the likelihood of ARGs under the SCAR model, we convert the ARG obtained from ARGweaver to the tskit tree sequence format [Kelleher et al., 2016]. The tskit tree sequence format provides a concise encoding of a set of local trees corresponding to the genealogy of the sample over different genomic regions [Kelleher et al., 2016, 2018]. The tree sequence uses tables of nodes and edges to define the topology of local trees within the ARG. The node table records the time and type of each event (e.g. sample, coalescent, recombination) in reverse chronological order going from tips to root. The edge table records the parent and child relationships between each node along with the coordinates [*left, right* [of the genomic region over which the parent-child relationship holds. This allows for a more efficient encoding of the local tree topologies, as edges present across multiple neighboring local trees can be compressed into a single edge with coordinates spanning multiple genomic regions. The tree sequence format also facilitates computing the likelihood of the ARG under the SCAR model. We can simply perform a post-order traversal through the ARG by iterating through the node table, computing the likelihood of the event at each node, update the edges (i.e. lineages) present in the ARG after each event, and computing the likelihood of no event occurring between nodes as in Equation (8). Code for converting ARGs into tskit tree sequence format and computing the likelihood of the ARG is available at https://github.com/sunnyfangfangguo/SCAR_project_repo.

### Simulation study

We simulated ARGs along with genomic sequence data to test the accuracy of ARG reconstruction using ARGweaver and the statistical performance of inference under the SCAR model before applying the method to real data. Simulated ARGs (tree sequences) were generated by msprime [Kelleher et al., 2016]. To test the accuracy of ARGweaver in reconstructing ARGs, sequence alignments for each local tree in an ARG were generated with a HKY substitution model [Hasegawa et al., 1985] with the transition/transversion ratio *κ* = 2.75 using Pyvolve [Spielman and Wilke, 2015]. Our simulations are similar to those of [Rasmussen et al., 2014b], which assumed a fixed effective population size *N*_*e*_ = 100, genome length *L* = 10, 000, and recombination rate per site per generation *r* = 2.5*e −* 06, and varied the mutation-to-recombination rate ratio *µ/r* from 1 to 2048. 100 simulations were conducted for each *µ/r* ratio.

Normalized Robinson-Foulds (RF) distances [Robinson and Foulds, 1981, Christensen et al., 2018] between corresponding simulated local trees and inferred local trees for each genome region were calculated as a metric of local tree accuracy, which varies between 0 and 1. RF distances along the whole chromosome were then calculated as an average RF distance over all genome regions. We also compared the true number of recombination events in the simulations to the number of recombination events inferred by ARGweaver. From Figure S1, we can clearly see that the ARG with the maximum iteration included the number of recombination events closest to the true number, while ARGs with the maximum likelihood consistently overestimated and ARGs with the maximum joint likelihood consistently underestimated the number of recombination events across all the ratios. Thus, we selected the ARG with the maximum iteration to show the accuracy of ARGweaver.

In order to test inference under the SCAR model, three simulation experiments were run: we (1) estimate the effective population size, recombination rate and migration rate directly from the true simulated ARG; (2) jointly estimate the recombination rate and migration rate from the true ARG; and (3) estimate the recombination rate from ARGs inferred by ARGweaver. When estimating migration rates between populations, we treat the ancestral location of each lineage as unknown and track the state of each lineage probabilistically. Again, 100 simulations were run for each simulation experiment. In the first two experiments, for each simulation, the true value of the estimated parameter(s) were drawn from a evenly spaced grid of values, while other parameters were kept constant. When only estimating the effective population size or recombination rate, we simulate ARGs without population structure.

## Results

### 1. Testing the accuracy of ARG inference using ARGweaver

#### Accuracy of local trees in ARGweaver inferred ARGs

Because our inference methods ultimately rely on the ability to accurately reconstruct ARGs, we first test the ability of ARGweaver to reconstruct ARGs from genomic data simulated under different mutation-to-recombination rate ratios *µ/r* in order to vary the number of phylogenetically informative sites (SNPs) between each recombination breakpoint. To evaluate the accuracy of ARGweaver, we use normalized Robinson-Foulds (RF) distances to quantify the topological differences between the simulated and reconstructed local trees in the ARG. From Figure 2 we can see that, with increasing *µ/r* ratios, the median RF distances decrease from 0.819 to 0.037, showing a clear increase in ARGweaver’s performance to accurately reconstruct the topology of local trees in the ARG. This is likely due to the fact that sequence diversity, and thus the number of phylogenetically informative sites between each recombination breakpoint, increases with *µ/r* ratio (Figure S2).

**Figure 2.**
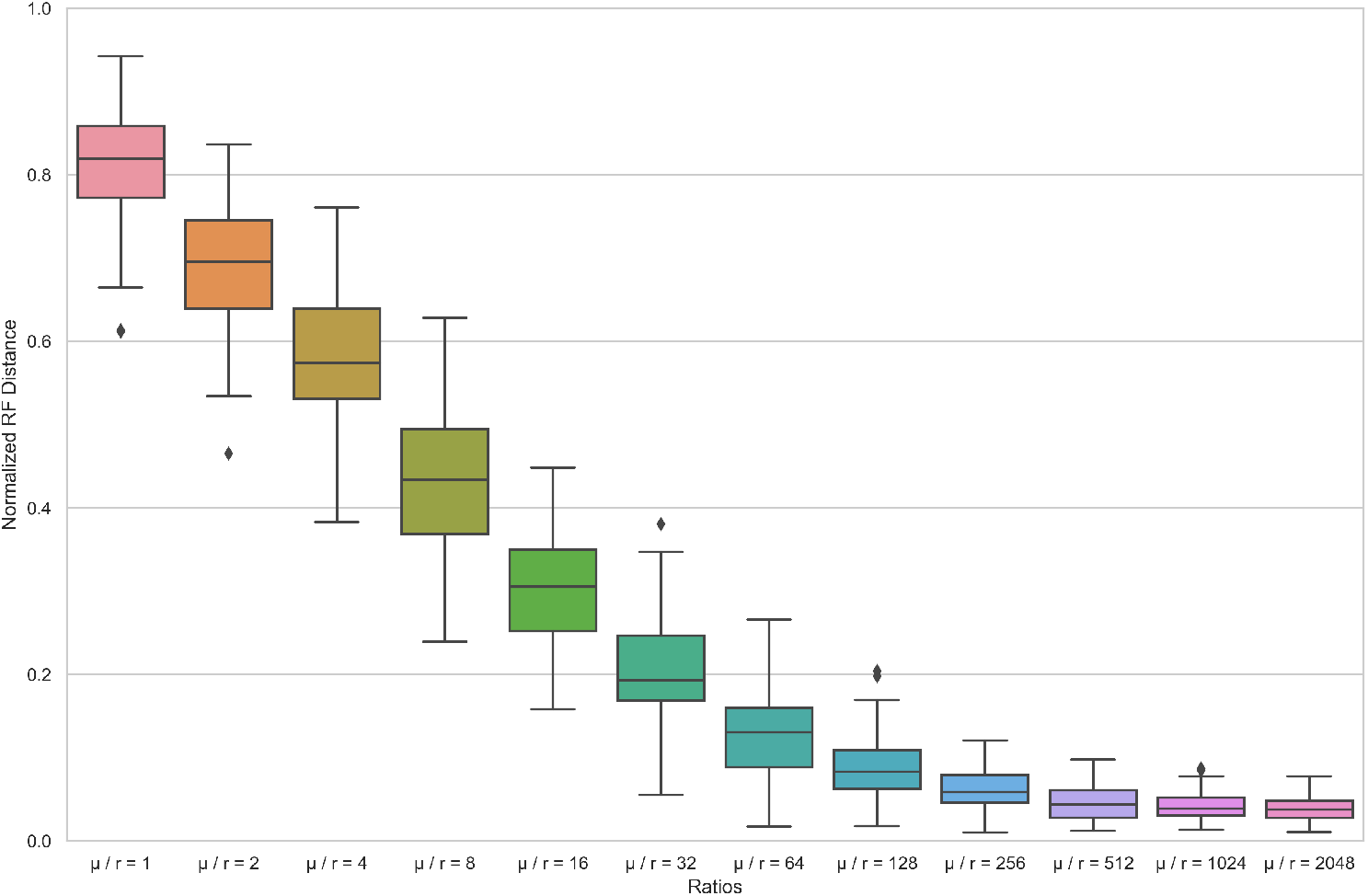
Normalized RF distances between the true (simulated) local trees and the local tree inferred by ARGweaver in the reconstructed ARG under different ratios of mutation rate / recombination rate. For each simulation, *N*_*e*_ is 100, sample size is 50, genome length is 1e04, and recombination rate *r* is 2.5e-06. Under each ratio, 100 simulations were run.

#### Accuracy in the number of inferred recombination events

We compared the number of recombination events inferred by ARGweaver against the true number known from simulations under 12 different *µ/r* ratios to further test the accuracy of ARGweaver. The number of recombination events inferred by ARGweaver was significantly and positively correlated with the true number of recombination events when the *µ/r* ratio ≥ 4, while at lower ratios the correlation is poor indicating it may not be possible to estimate the true number of recombination events unless the mutation rates is at least several times higher than the recombination rate (Figure 3). As the *µ/r* ratio increases, the correlation generally becomes stronger.

**Figure 3.**
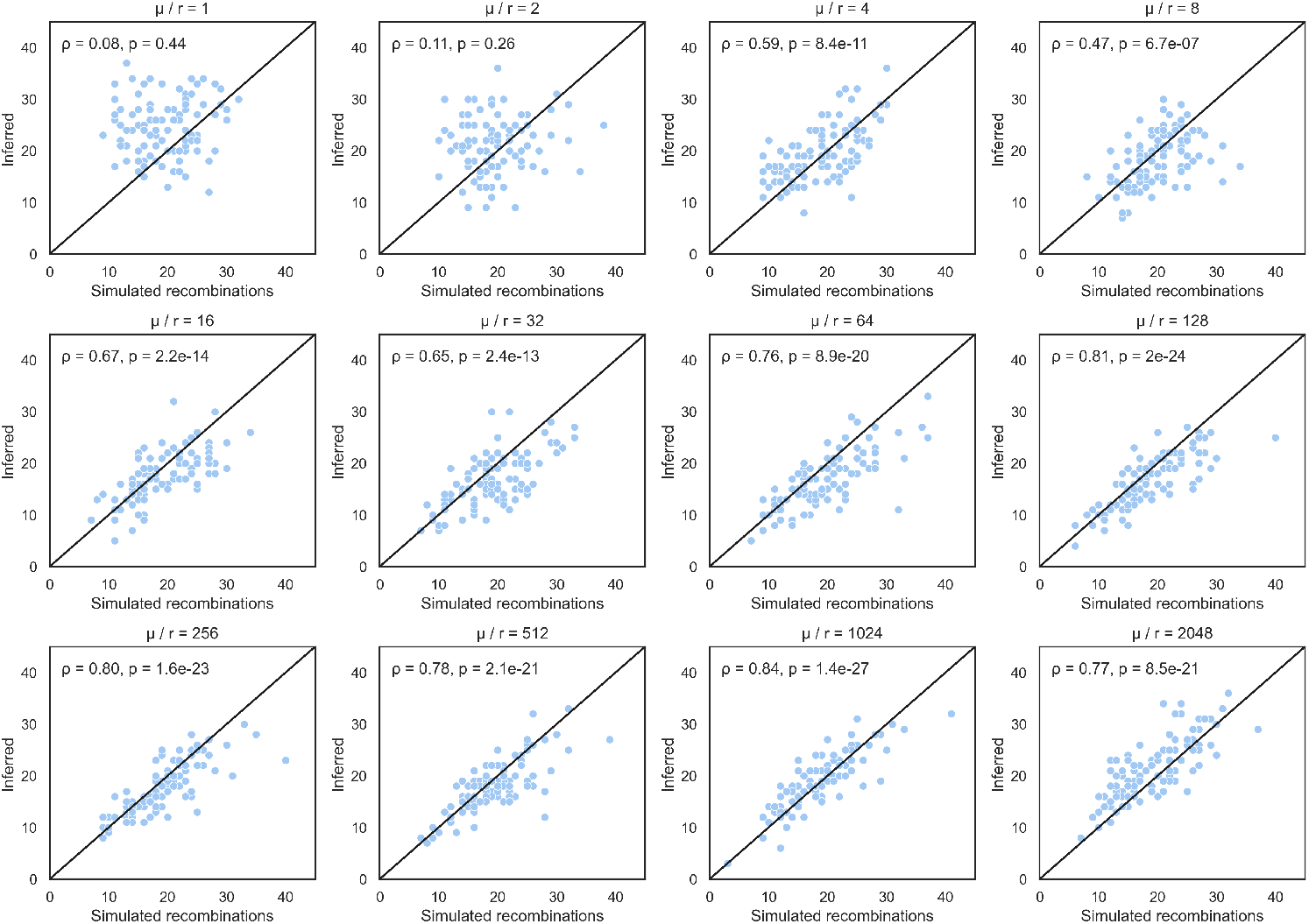
Simulated number of recombination events versus number inferred by ARGweaver under different ratios of mutation rate / recombination rate. Each diagonal line is *x* = *y. ρ* and *p* are the correlation coefficient between the simulated and estimated number of events and corresponding p-value, respectively. We selected the ARG sampled by ARGweaver during the final MCMC iteration to count the number of recombination events.

### 2. Testing the SCAR model on known ARGs

#### Statistical performance of estimating *N*_*e*_, recombination rate *r*, and migration rate *M*

Next, we tested how well we are able to estimate effective population sizes *N*_*e*_, recombination rates *r*, and migration rates *M* from simulated ARGs known without error under the SCAR model. Figure 4 shows that the SCAR model can accurately estimate all three of these parameters across a wide range of true values. Table 1 summarizes the performance of our estimates across simulations in terms of the relative bias, coverage of the 95% credible intervals, and calibration between true and estimated parameters. We find that migration rate estimates are very accurate when the true migration rates are smaller than 1 per unit time. However, in some simulations the migration rates are overestimated, especially when the true rate was larger than 1, indicating an inability to precisely estimate high rates likely due to the fact that the likelihood function becomes very flat across a wide range of higher rates. After testing the SCAR model on simulations with different sample sizes (Figure S3), we found that the additional information provided by increased sampling could provide more accurate and precise migration rate estimates.

**Table 1.** The relative bias, coverage, and calibration of estimating *N*_*e*_, *r, M* by the SCAR model

| Parameter | Relative error | Coverage | Calibration |
| --- | --- | --- | --- |
| $N_e$ | +1.23% | 92 in 100 times | 0.98 |
| $r$ | +1.87% | 97 in 100 times | 0.98 |
| $M$ | +22.36% | 95 in 100 times | 0.85 |

**Figure 4.**
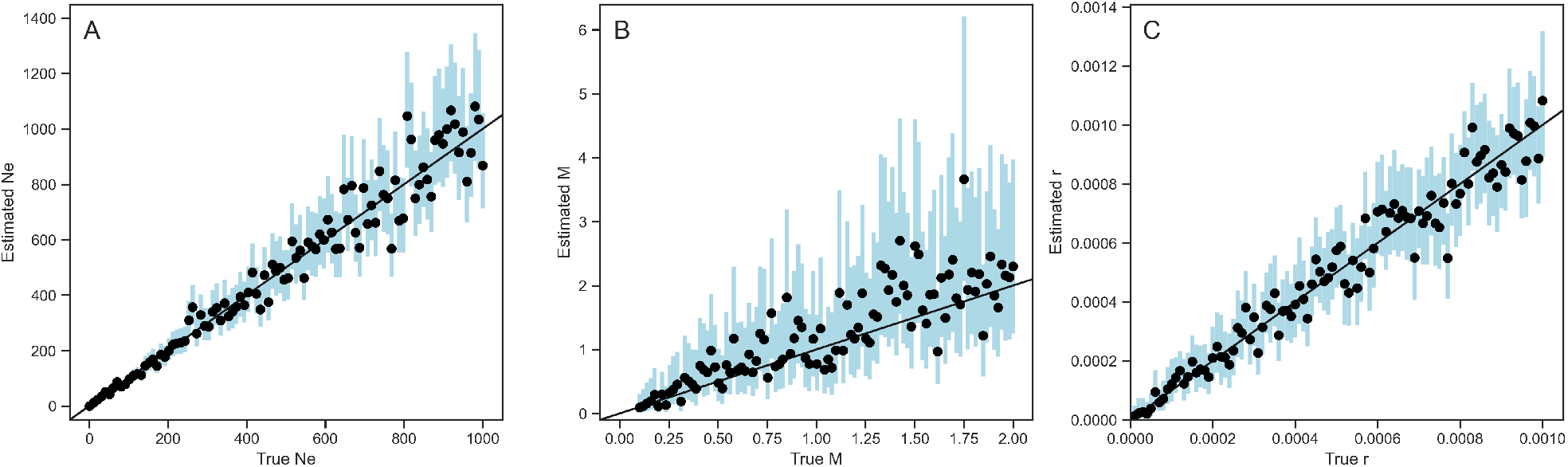
Inference of effective population size *N*_*e*_ (A), migration rate *M* (B), and recombination rate *r* (C) from 100 known ARGs. Migrations rates are assumed to be equal (symmetric) between two populations. Dots and blue bars represent the median posterior estimates and the 95% confidence intervals for each simulation.

We further tested the performance of the SCAR model when jointly estimating the recombination rate *r* and migration rate *M* together. As shown in Figure 5, the SCAR model can accurately infer the marginal posterior distribution of each parameter even when the two parameters are jointly estimated together.

**Figure 5.**
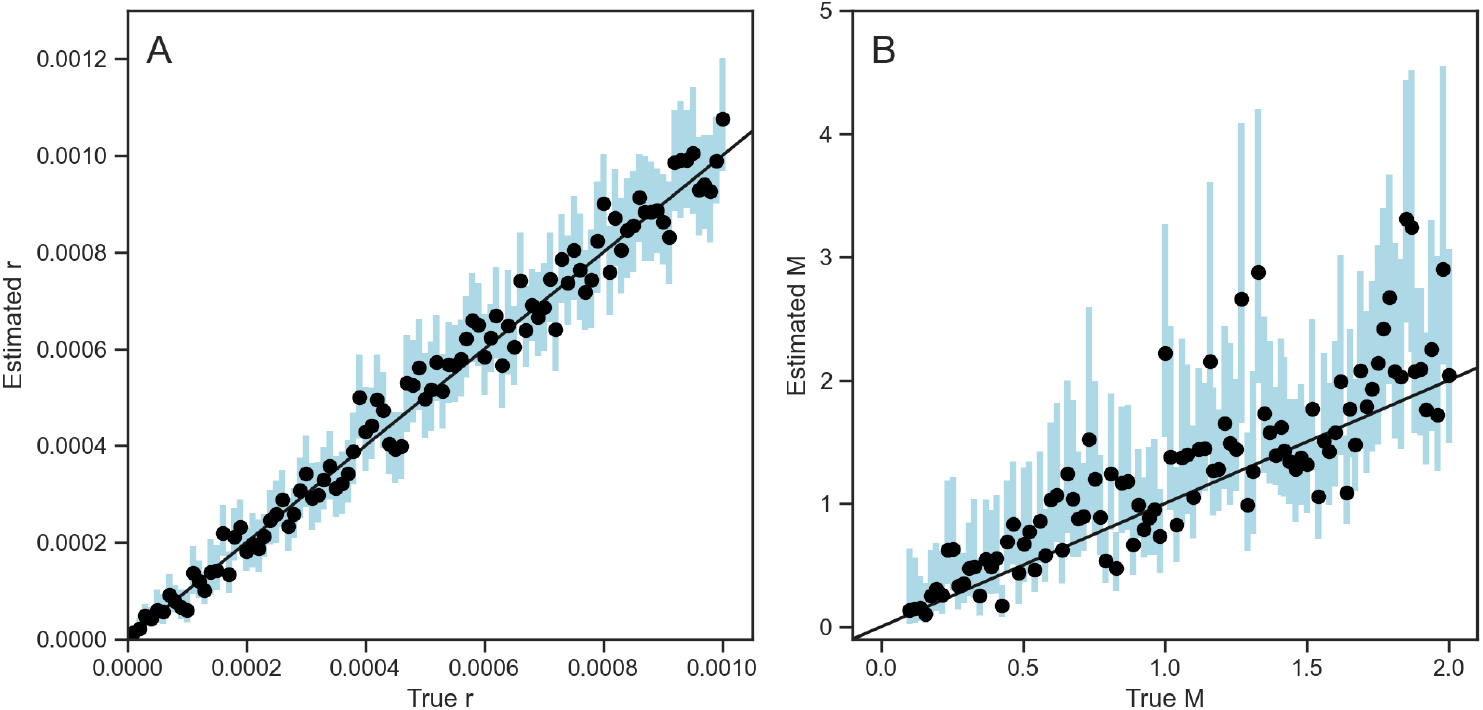
Joint estimation of recombination and migration rates together. Dots and blue bars represent the median posterior estimates and the 95% confidence intervals of the marginal posterior distribution of each parameter from each simulation.

#### Statistical performance of estimating recombination rates from ARGs inferred by ARG-weaver

In order to see how ARG reconstruction errors influence our estimates, we estimated recombination rates using the SCAR model from ARGs reconstructed by ARGweaver rather than the true simulated ARGs. We also compared to the recombination rates estimated by ARGweaver, which simply counts recombination events in the reconstructed ARG and divides by the total branch-length of the ARG to estimate the recombination rate [Hubisz and Siepel, 2020]. From Figure 6 we can see that the accuracy of recombination rates estimated by both SCAR and ARGweaver improves with increasing *µ/r* ratios. However, when the *µ/r* ratio exceeds 1024, recombination rates become slightly over-estimated. We suspect that the increased accuracy of recombination rate estimates at higher *µ/r* ratios is due to increasing phylogenetic information about the local trees and thus power to distinguish true recombination events from uncertainty in the topology of local trees. However, at very large ratios individual sites may become phylogenetically uninformative due to recurrent or convergent mutations (i.e. saturation effects) and we therefore may become overconfident that discordance between local trees is due to recombination rather than phylogenetic errors. Compared with ARGweaver, SCAR can estimate the recombination rate with less variance between simulations.

**Figure 6.**
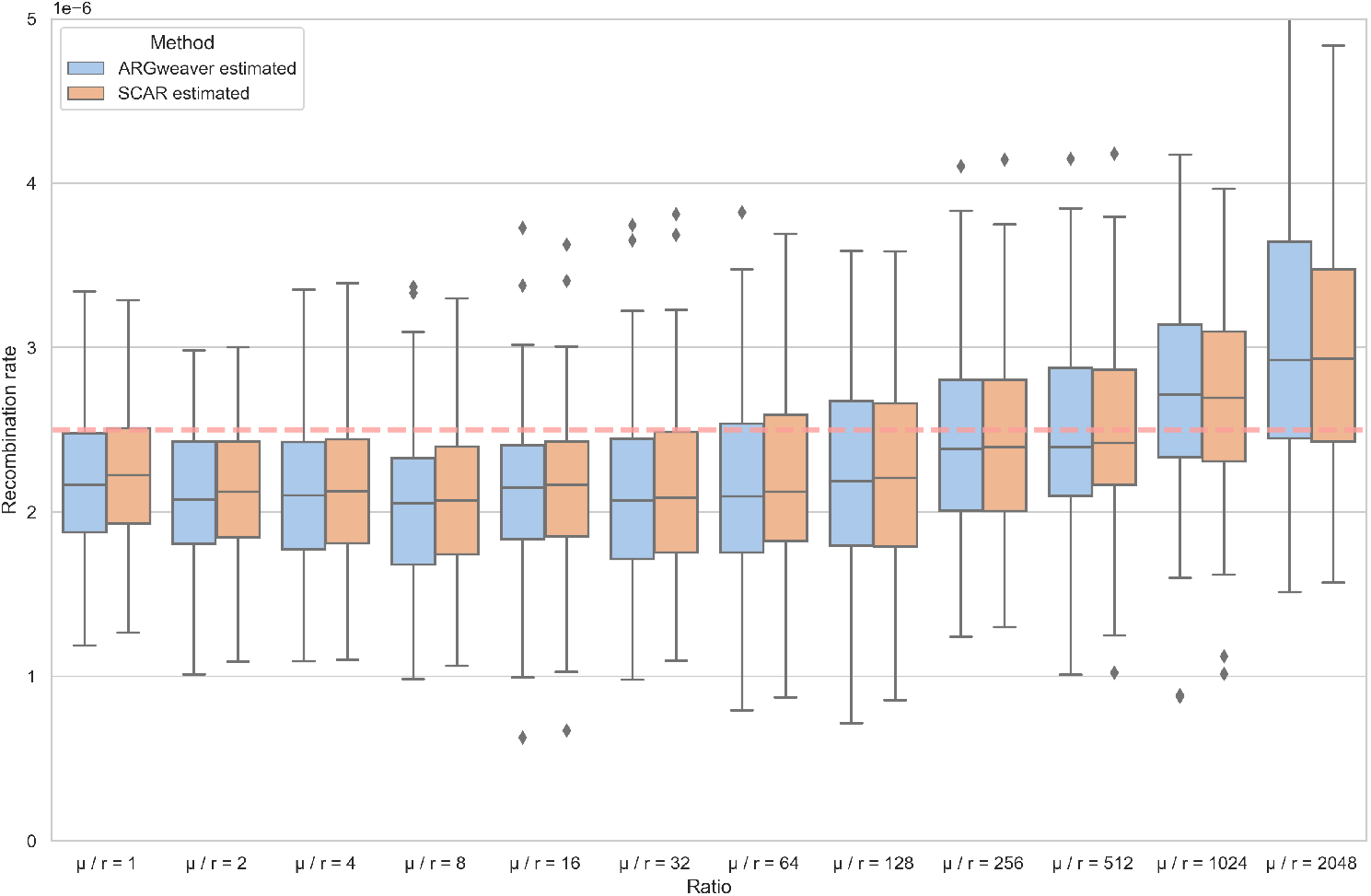
Recombination rates estimated using ARGweaver and SCAR from ARGs inferred by ARGweaver under different *µ/r* ratios. The dashed red line is the simulated recombination rate in all simulations. Under each *µ/r* ratio, 100 simulations were run.

### 3. Recombination and migration in *Aspergillus flavus*

#### Introduction

It is estimated that over 25% of food crops are contaminated with mycotoxins worldwide [Eskola et al., 2020]. *Aspergillus flavus*, a pathogen of plants, animals and insects, is a major aflatoxin producer that has a broad economic impact [Klich, 2007]. *A. flavus* can infect and contaminate preharvest and postharvest seed crops with the carcinogenic secondary metabolite aflatoxin [Amaike and Keller, 2011]. This fungus is predominantly haploid and homokaryotic [Runa et al., 2015]. *A. flavus* was thought to be cosmopolitan and clonal, until evidence for genetic recombination due to a cryptic sexual state were reported [Geiser et al., 1998] and later the sexual stage was described [Horn et al., 2009]. In natural populations, *A. flavus* undergoes both sexual and asexual reproduction [Horn et al., 2016, Ojeda-Lopez et al., 2018]. Previous studies also found extensive recombination in the ancestral history of the aflatoxin cluster [Moore et al., 2009, 2013], which is a 70-Kb-gene-cluster near chromosome 3’s right telomeric region [Amaike and Keller, 2011, Carbone et al., 2007]. Based on multilocus DNA sequence markers in the aflatoxin cluster (*aflM/aflN* and *aflW/aflX*) and three other nuclear loci (*mfs, amdS, trpC*), this fungus can be delimited into two evolutionary distinct lineages: IB and IC, where IB includes mainly nonaflatoxigenic isolates while IC includes both toxigenic and atoxigenic strains [Moore et al., 2009, 2017]. There is evidence that *A. flavus* has the potential for long-distance dispersal via conidia [Wicklow et al., 1993, Probst et al., 2011, Ortega-Beltran et al., 2020], but movement between geographic locations in poorly characterized.

Given that both recombination and migration shape the evolution history of *A. flavus*, here we aim to use ARGweaver and our SCAR model to explore the two evolutionary forces together by reconstructing ARGs, and then estimating the recombination and migration rates.

#### Materials and Methods

The genome size of *A. flavus* is about 37 Mb on eight chromosomes [Fountain et al., 2020], but we focused our analysis on the migration and recombination history of chromosome 3, which is about 5 Mb. A total of 51 lineage IB strains and 48 lineage IC strains were collected across the United States, including Arkansas, Indiana, North Carolina, and Texas in 2013 [Molo et al., 2022]. Sample metadata is provided in supplementary file *sample name population*.*xlsx*. Single-nucleotide polymorphism (SNP) genotyping was performed across chromosome 3 with *A. oryzae* RIB40 as the reference genome [Machida et al., 2005]. Because there was limited migration between lineages IB and IC [Molo et al., 2022], we analyzed the two lineages separately. No SNPs in the aflatoxin gene cluster were included for IB because few isolates harbored this gene cluster.

We used ARGweaver to infer ARGs from SNPs spanning most of chromosome 3. Because ARGweaver requires an estimate of the recombination rate to infer ARGs, we used LDhat version 2.2 [McVean et al., 2002] to estimate Watterson’s theta and the population recombination rate. SNP data was filtered using a series of different missing data thresholds before running LDhat, and we estimated the median recombination rate across filtered data sets. The *A. flavus* mutation rate was previously estimated as 4.2e-11 per site per mitosis [Álvarez Escribano et al., 2019], which can be converted to 2.82e-09 per base per generation. Given this mutation rate, the effective population size *N*_*e*_ was calculated from Watterson’s theta. ARGweaver also needs a maximum time threshold for coalescent events, which was set as the expected time to the most recent common ancestor based on the sample size and estimated population sizes. With these parameters, we ran ARGweaver for 20,000 iterations with 1000 iterations as burn-in. To keep ARGweaver’s run time manageable, we compressed blocks of 5 variable sites by conditioning the breakpoints between each block in a flexible manner so that no more than one variant site in the same block was chosen [Hubisz and Siepel, 2020]. All the runtime parameters can be found in the supplementary file (*ARGweaver parameters*.*xlsx*). From the SMC files produced in the iterations, we choose the ARG with the maximum joint-likelihood as our best estimate of recombination patterns across chromosome 3.

We used a tanglegram to show the topological changes between neighboring local trees in the inferred ARG. In a tanglegram, each local tree is drawn and then auxiliary lines are drawn to connect matching taxa in neighboring trees. If there is no recombination, the lines connecting matching taxa should be horizontal whereas crossing lines can be used as a visual heuristic to assess the extent of recombination. We use the python package baltic [Dudas et al., 2016] to display the tanglegrams. The ARGs reconstructed in ARGweaver were then used to estimate recombination and migration rates using our SCAR model. For these analyses we used exponential priors (for IB *r* ∼ *Exp*(1.37*e −* 09), *m*_*ij*_ ∼ *Exp*(0.1); for IC *r* ∼ *Exp*(2.17*e −* 10), *m*_*ij*_ ∼ *Exp*(0.1)). MCMC chains were run for 40,000 iterations. Besides estimating these parameters from the reconstructed ARGs, we also compared the posterior distributions of estimated migration rates from the consensus tree of all the local trees in the ARG along the whole chromosome using the SCAR model. Additionally, we compared how the RF distances between pairs of local trees for different regions of the genome changed based on their genomic distance, which was the absolute value of coordinates (middle of genome segment location) of the difference between two trees.

## Results

### 1. The reconstructed ARGs for chromosome 3 of lineages IB and IC

The *µ/r* ratios were calculated using Watterson’s *θ* and the population recombination rate obtained by LDhat. For lineage IC, the *µ/r* ratio of the entire chromosome 3 was 13, whereas for lineage IB *µ/r* was 2.05. Even though the *µ/r* ratio of lineage IB was slightly lower than the lower limit at which we found ARGweaver could accurately reconstruct ARGs in simulations, we continued with the analysis in order to explore the limits of ARG-based phylogeographic inference.

We reconstructed ARGs for chromosome 3 of the *A. flavus* genome from 51 lineage IB isolates and 48 lineage IC isolates. Overall, the ARGs contained 190 recombination events for lineage IB and 774 recombination events for lineage IC. To visualize how the topology of local trees varied across the genome, we plotted tanglegrams for the first 10 local trees in each ARG for lineage IB (Figure 7A) and lineage IC (Figure 7B), as well as the 12 local trees in the aflatoxin gene cluster for lineage IC (Figure 7C). Although there was always one recombination event between each local tree in the ARG reconstructed by ARGweaver, not all recombination events result in topological discordance between neighboring trees. In the ARG of lineage IB, 92.1% of recombination events caused topological discordance between local trees whereas the other 7.9% only changed coalescent times. In the ARG of lineage IC, 98.3% of recombination events resulted in topological discordance while only 1.7% caused changes in coalescent times. The average normalized RF distance between local trees for IB and IC was 0.094 and 0.115, respectively (Figure 8A), indicating considerable phylogenetic discordance between local trees. However, some phylogenetic discordance may be due to errors in reconstructing local trees, especially because the genetic distance between pairs of breakpoints were often relatively small (Figure S4A), such that many non-recombining segments likely did not have enough segregating sites to reconstruct local tree accurately. Overall though, we found that the normalized RF distance between pairs of local trees increased logarithmically with their distance from each other in the genome (Figure S4B), consistent with discordance being driven by recombination rather than phylogenetic error over larger genomic distances.

**Figure 7.**
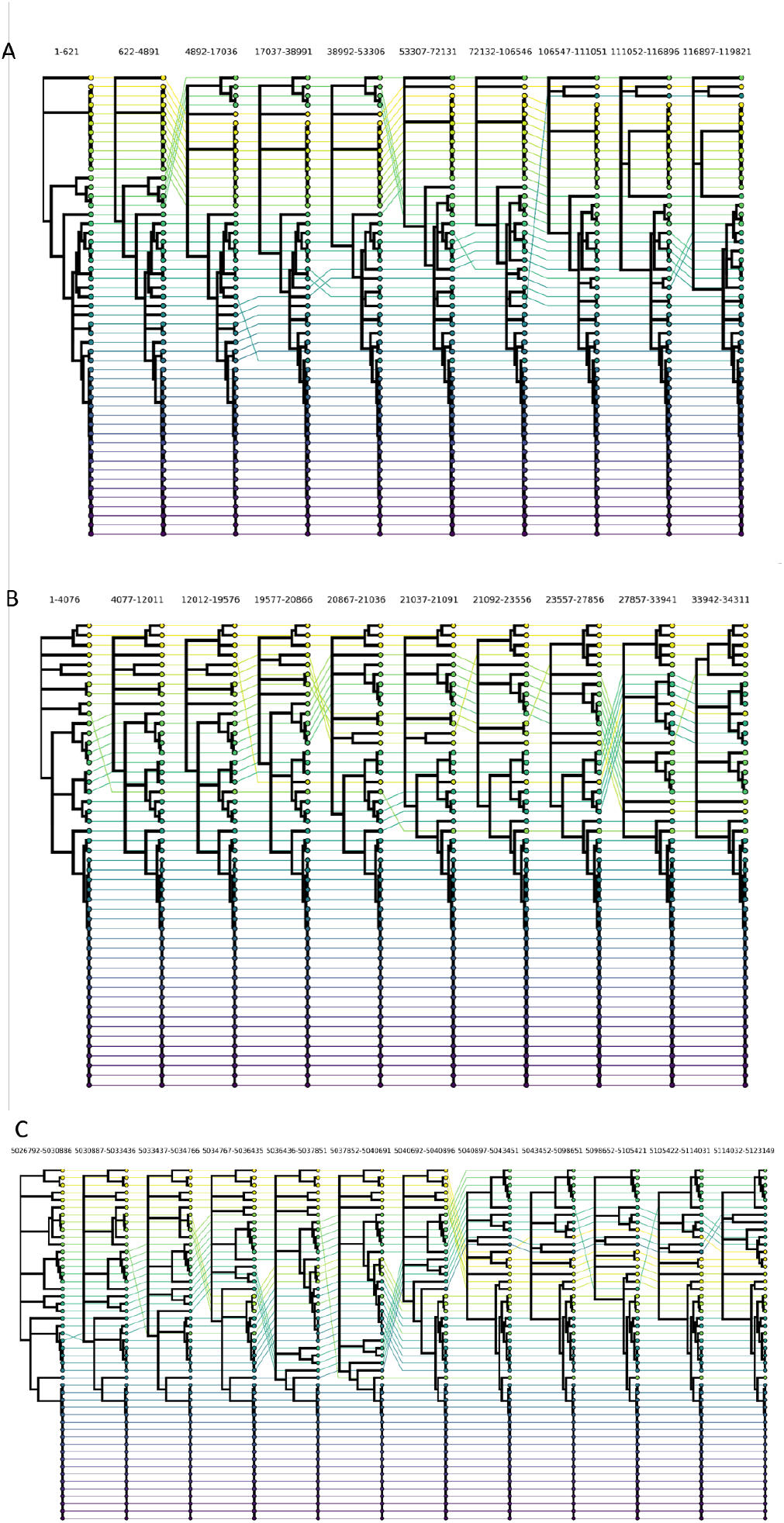
ARG of the 51 lineage IB isolates (A), 48 lineage IC isolates (B) and the aflatoxin gene cluster of lineage IC (C) reconstructed by ARGweaver. The reconstructed ARG is visualized using a tanglegram to show how the topology of local trees varies across chromosome 3. Each local tree corresponds to one genome region separated from neighboring regions by an inferred recombination breakpoint. Note only the first 10 of 193 local trees in the ARG of lineage IB, and only the first 10 of 775 local trees in the ARG of lineage IC are shown. In the ARG of the aflatoxin gene cluster, there are 12 local trees.

**Figure 8.**
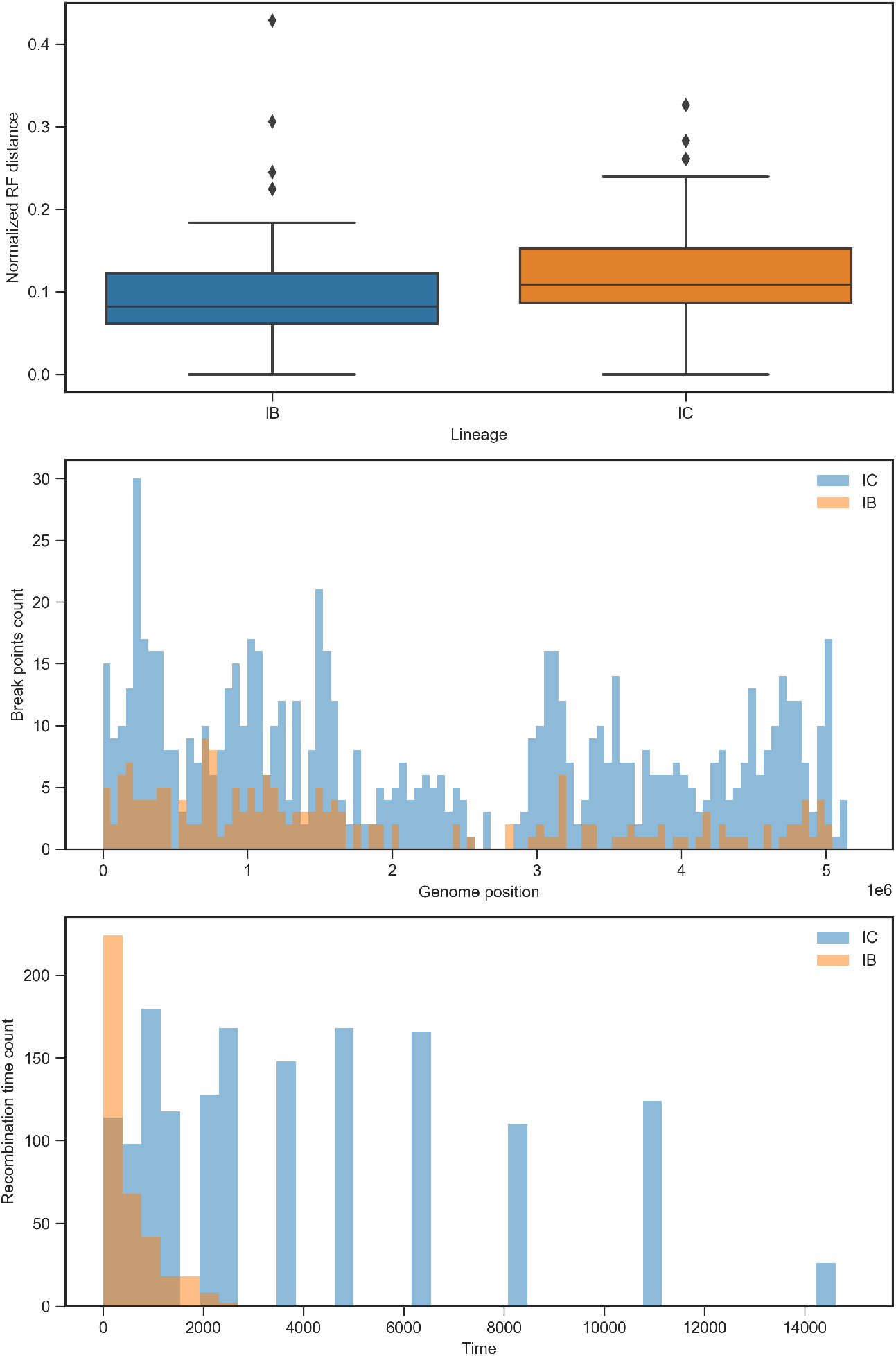
Normalized RF distance between neighboring local trees (A), the frequency of recombination events distribution along the genome (B), and the frequency of recombination time (generations in the past) of lineages IB and IC (C), respectively.

Recombination breakpoints were distributed unevenly across the genome, with the centromeric region containing far less recombination events for both lineages (Figure 8B). Figure 8C shows the distribution of recombination times for both lineages. While recombination events occurred mostly in the recent past for lineage IB, many recombination events occurred in the much deeper past for lineage IC.

### 2. Recombination rate of lineages IB and IC

Using the SCAR model, we estimated the recombination rate for lineages IB and IC (first column of Figure 9). The recombination rate of lineage IB was estimated to be 2.28E-09 per site per generation, which is about 0.81X lower than the mutation rate, which we assume is constant across both lineages and all of chromosome 3. The recombination rate of lineage IC was estimated to be 1.06E-09 per site per generation, which is about 0.38X lower than the mutation rate. Although less recombination events were identified for lineage IB than lineage IC, lineage IB was estimated to have a higher recombination rate. This counter-intuitive result can be explained by the fact that lineage IB also has a smaller effective population size and thus coalescent times occur in the more recent past, resulting in less time for recombination events to occur in IB than in IC, consistent with the temporal distribution of recombination events observed in Figure 8C.

**Figure 9.**
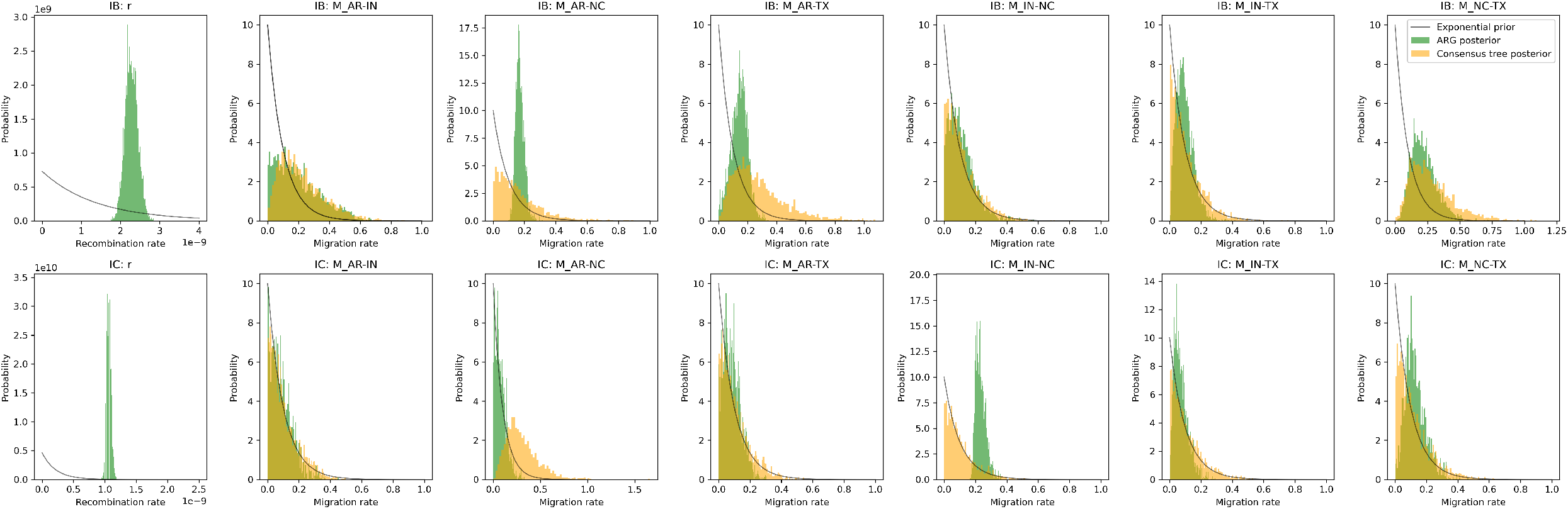
The prior and posterior distributions of demographic parameters estimated from either the full ARG or a single consensus tree for lineages IB and IC. For lineage IB, the mean value of exponential prior of recombination rate is 1.37e-09, and the mean value of exponential prior of migration rates is 0.1; For lineage IC, the mean value of exponential prior of recombination rate is 2.17e-10, and the mean value of exponential prior of migration rates is 0.1. Note that we cannot estimate *r* from the consensus tree so only the posterior distribution estimated from the ARG is shown.

### 3. Migration rates of lineages IB and IC between subpopulations

Using the SCAR model, we estimated the migration rates of lineages IB and IC between subpopulations along with their recombination rate from their ARGs (Figure 9). Migration rates between subpopulations were found to vary between 0.05 and 0.2 migrations per generation, suggestive of extensive movement between populations. Migration rates between subpopulations were similar within each lineage (Figure 10). However, for lineage IB, the migration rate between subpopulations in North Carolina and Texas was slightly higher, and the Arkansas subpopulation had the highest migration rates to other subpopulations overall; for lineage IC, the migration rate between subpopulations in Indiana and North Carolina was slightly higher, as well as the migration rate between subpopulations in Texas and North Carolina. Generally, the migration rates of lineage IC were lower than for lineage IB.

**Figure 10.**
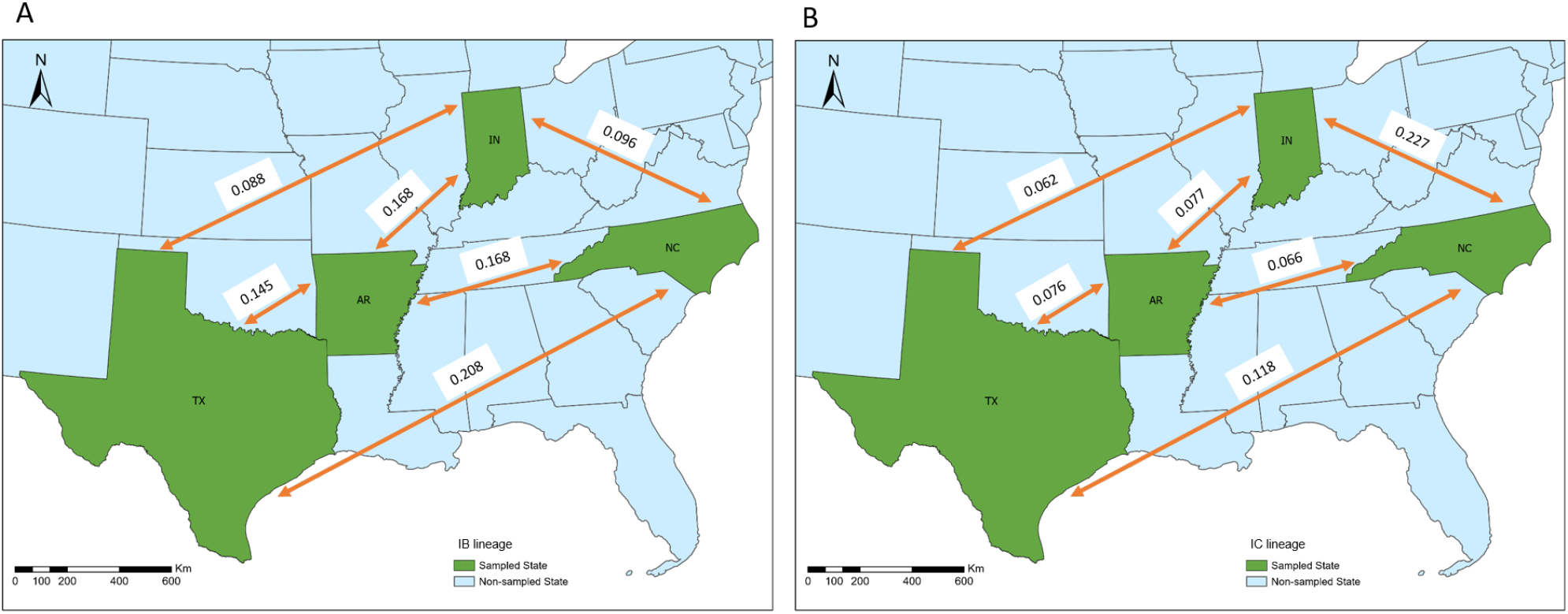
The migration rates (per generation) estimated for lineages IB (A) and IC (B) between subpopulations in four states.

We also compared migration rates estimated from full ARGs against migration rates estimated from a single phylogeny, in this case the consensus tree of each ARG. The posterior distribution of migration rates inferred from both the full ARG and the consensus tree are compared in Figure 9 and provided in supplementary file *summary statistics arg consensustree*.*xlsx*. Overall, the posterior distributions of migration rates estimated from the consensus tree diverged little from the prior distribution, indicating that the consensus trees contained little information about migration patterns. By contrast, the posterior distributions estimated from the full ARG were typically peaked with a much greater probability density concentrated around the posterior median relative to the prior distribution. These results suggest that we can obtain much more information from the full ARG than from any individual consensus tree (or gene tree), owing to the greater number of ancestral lineages and their associated migration histories in the ARG.

## Discussion

Because population structure and recombination jointly shape the genealogical history of many organisms, we developed SCAR to extend the structured coalescent to include ancestral recombination. When used for demographic inference, we showed that SCAR can successfully estimate effective population sizes, migration rates and recombination rates from reconstructed ARGs. We then showed that SCAR can recover these parameters accurately both from the true (simulated) ARGs and from ARGs reconstructed from genomic sequence data using ARGweaver, although performance declines as the recombination rate approaches the mutation rate. We also applied the SCAR model to *A. flavus* genomic data using ARGs inferred by ARGweaver, demonstrating how these methods can be applied to real world pathogens with complex histories of both migration and recombination.

While new methods for ARG reconstruction are being developed at a rapid pace, we chose ARGweaver as a gold-standard for inference as it reconstructs ARGs by sampling them from their full posterior distribution up to the approximation introduced by the Sequential Markov Coalescent [McVean and Cardin, 2005, Rasmussen et al., 2014b], which is known to be a very good approximation to the full coalescent with recombination [Wilton et al., 2015]. Other, more approximate methods, may therefore be faster or allow larger samples sizes but are unlikely to outperform ARGweaver in terms of accuracy. We therefore used ARGweaver to explore the limits of reconstructing ARGs and estimating recombination rates from simulated data. Regardless of method, the number of phylogenetically informative sites (i.e. SNPs) between recombination breakpoints is likely the ultimate factor limiting accurate reconstruction of local tree topologies within an ARG and thereby our ability to distinguish true recombination events from topological discordance introduced by phylogenetic uncertainty. We therefore explored the limits of accurate ARG reconstruction by varying the ratio of the mutation rate to the recombination rate *µ/r*. We found that at high *µ/r* ratios, ARGweaver does in fact reconstruct ARGs very accurately. However our ability to reconstruct local trees within the ARG rapidly degrades at lower *µ/r* ratios and, as expected, our ability to accurately estimate recombination rates likewise decreases with our ability to accurately reconstruct ARGs. Our simulations suggest that a *µ/r* ratio of about 4 poses a practical lower limit on our ability to reconstruct ARGs. While many rapidly evolving viruses and predominately clonal bacteria exceed this threshold [Awadalla, 2003, Vos and Didelot, 2009], this definitely poses a challenge to accurate ARG reconstruction for many highly-recombining bacteria and eukaryotic organisms. For example, *µ/r* ratios for fungi have been reported as low as 0.1 in *Zymoseptoria triciti* [Stukenbrock and Dutheil, 2018] *and as high as 45*.*9 for Glomus etunicatum* [den Bakker et al., 2010]. However, because recombination requires direct physical interactions (e.g. sexual reproduction), recombination rates can vary substantially even between populations of the same species based on the frequency at which individuals encounter one another [Hasan and Ness, 2020, Stapley et al., 2017]. This suggests that ARG reconstruction methods will likely need to be applied on a one-by-one basis to particular data sets rather than being applied or dismissed for broad classes of organisms.

The SCAR model tracks the movement of lineages in an ARG between subpopulations by approximating ancestral state probabilities. Rather than jointly estimating the ancestral states with the other demographic parameters, we probabilistically track the movement of lineages and integrate over their unknown states using the approach first developed by Volz [2012]. Using this method, we can accurately and quickly estimate migration rates from simulated ARGs. However, we found that SCAR overestimates migration events to some extent, especially when the true migration rates approach one per generation. This bias might be caused by assuming lineage state probabilities evolve independently across lineages and are independent of the coalescent process [Volz, 2012]. However, Müller et al. [2017] showed that the lineage independence assumption performs worst when migrations rates are very low relative to coalescent rates (the opposite of our situation) and when coalescent rates are highly asymmetric between populations (a situation we do not consider). We therefore think it more likely that, for larger migration rates, there simply is not enough information to determine the true rate, as the likelihood surface remains essentially flat across a wide range of higher values (Figure S5). Based on the results of estimating migration rates using different sample sizes, we show that the larger the sample size, the more accurate estimation becomes, verifying our speculation that biases in estimating higher migration rates were attributable to a lack of information.

Several earlier approaches likewise aimed to extend the structured coalescent to include recombination, including LAMARC 2.0 [Kuhner, 2006], CSD 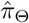 [Steinrücken et al., 2013], ARGweaver-D [Hubisz et al., 2020], and SCoRe [Stolz et al., 2022]. LAMARC 2.0 can simultaneously estimate migration rates, population growth rates, and recombination rates [Kuhner, 2006]. SCAR and LAMARC 2.0 model the recombination process in the same way Kuhner et al. [2000], but like other early implementations of the structured coalescent [Beerli and Felsenstein, 2001], it uses MCMC to sample migration histories, limiting its applicability to larger data sets or data sets with more than a few sampled populations [Kuhner, 2009]. ARGweaver-D [Hubisz et al., 2020] is a major extension of ARGweaver and allows a user to define their own demographic model and infer migration events between an arbitrary number of subpopulations. Finally, SCoRe [Stolz et al., 2022] can infer migration rates and reassortment patterns for segmented viruses from a phylogenetic network jointly estimated in BEAST2 [Muller et al., 2020]. Conceptually, SCoRe is very similar to SCAR in that both methods track the movement of lineages probabilistically based on similar approximations to the structured coalescent, although SCoRe allows for more refined approximations than currently implemented in SCAR. The main difference between SCAR and SCoRe is that the SCoRe model specifically focuses on reassortment, where different segments of a viral genome are inherited from different parents, leading to a block-like haplotype structure where all sites in the same segment necessarily share the same phylogenetic history. In contrast, SCAR allows for a more general model of recombination where recombination breakpoints and thus changes in local tree topologies can occur anywhere across the genome, leading to much more complex ARGs. The SCAR model therefore accommodates varying mechanisms of recombination, such as crossovers and gene conversion, and so applicable to a broader range of viral, bacterial and fungal genomes.

For organisms like *A. flavus* that recombine frequently relative to their mutation rate, there may be further challenges to inferring ARGs given a low *µ/r* ratio. Furthermore, the aflatoxin gene cluster is reported to be a recombination hot spot [Moore et al., 2009], so the *µ/r* ratios likely vary across the genome, but we assume a constant mutation rate and recombination rate prior when reconstructing ARGs. While there are likely regions where recombination rates are lower and we can accurately reconstruct phylogenetic relationships, this will not be the case across the entire genome. Despite the inherent variability in ARG reconstruction accuracy across the genome, our estimates using ARGweaver/SCAR are consistent with those reported in the *A. flavus* literature. We found that the ratio of the recombination rate to mutation rate of lineages IB and IC was 0.81 and 0.38, respectively. In study of Drott et al. [2020], the recombination to mutation ratio in three populations calculated by ClonalframeML are from 0.19 to 0.44. Even though our calculations were based on a single chromosome, they were similar in magnitude to genome-wide estimates for lineage IC. Moreover, we found that the centromeric region of the genome contain far less recombination events for both lineages IB and IC, which accords with the knowledge that regions surrounding centromeres are a cold spot of recombination [Choo, 1998, Stapley et al., 2017].

While incorporating recombination into phylogeography has typically been viewed as burdensome, considering recombination and the full ancestry of sampled genomes through an ARG allows us to track the ancestral movement of many different genes or genomic regions. Considering the full ARG rather than just a single phylogeny therefore provides more information about demographic parameters and allows us to see how migration histories vary across the genome. While it has long been appreciated that considering multiple ancestral histories across the genome can improve demographic inference [Wakeley and Hey, 1997, Hare, 2001, Goss, 2015], here we demonstrate that reconstructing ARGs for *A. flavus* provides much more information about migration between populations than does a single (consensus) tree. Indeed, posterior distributions for the *A. flavus* migration rates inferred from ARGs are concentrated around their posterior median while the same migration rates inferred from a single tree diverge little from the prior, demonstrating that it may be possible to estimate migration rates from ARGs even when a single tree contains no information about these parameters.

Using the SCAR model, we can now conduct phylogeographic inference using all the information contained within an ARG. In the future, we plan to combine the SCAR model with more computational efficient methods for reconstructing ARGs like Espalier [Rasmussen and Guo, 2022] to explore how recombination and migration jointly shape the phylogeographic history of a broad range of pathogens.

## Acknowledgments

This research was supported by grant 2019-67021-29932 from the USDA NIFA. DAR was additionally supported by funding from USDA Hatch project 1016556.

## Supplementary Materials

**Figure S1.**
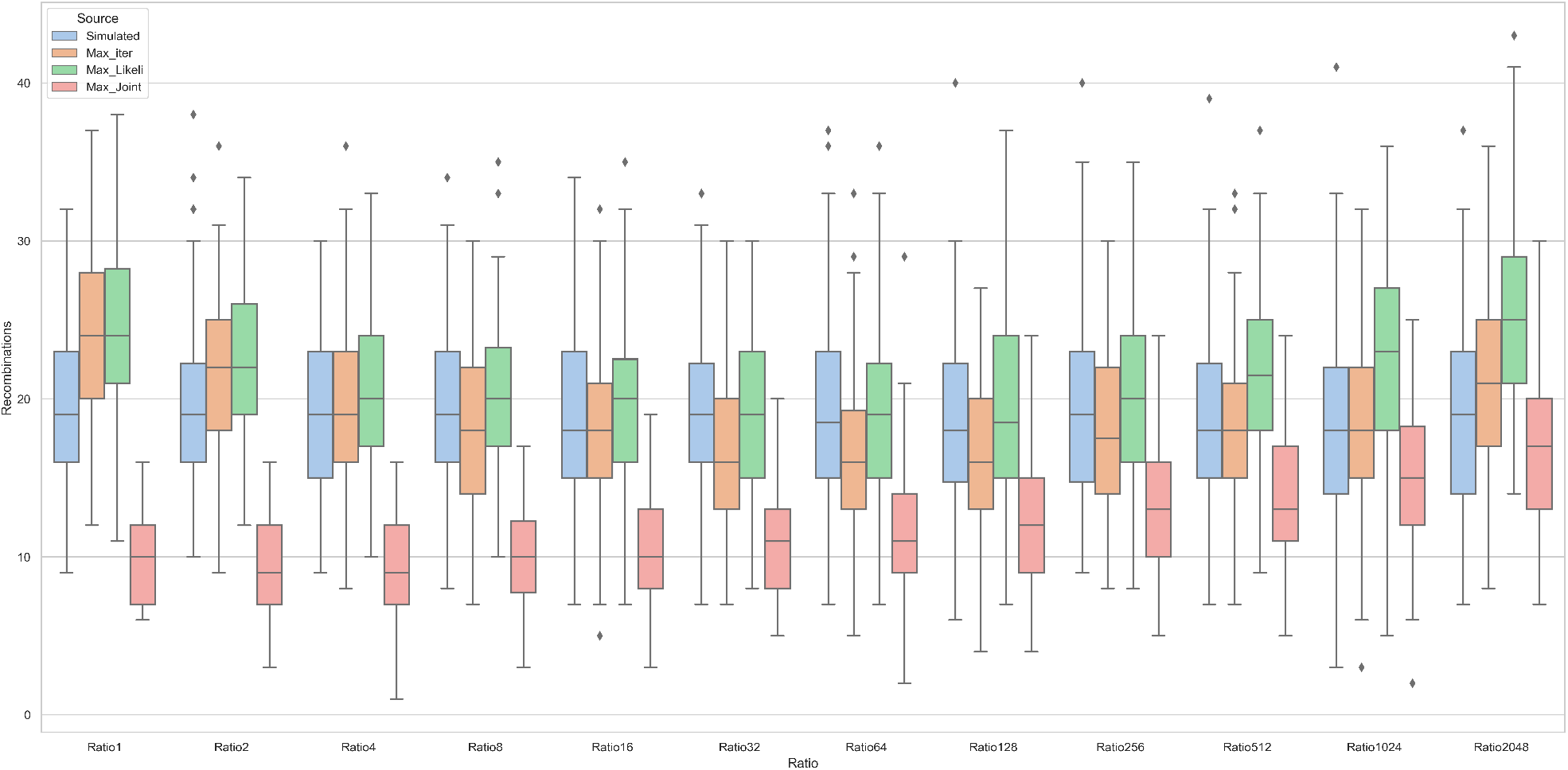
The SMC files with the maximum joint likelihood, maximum likelihood, and maximum iteration inferred numbers of recombination events compared with the true simulated numbers under different *µ/r* ratios. In the legend, *Max iter, Max Likeli, Max Joint* represents maximum iteration, maximum likelihood, and maximum joint likelihood, respectively

**Figure S2.**
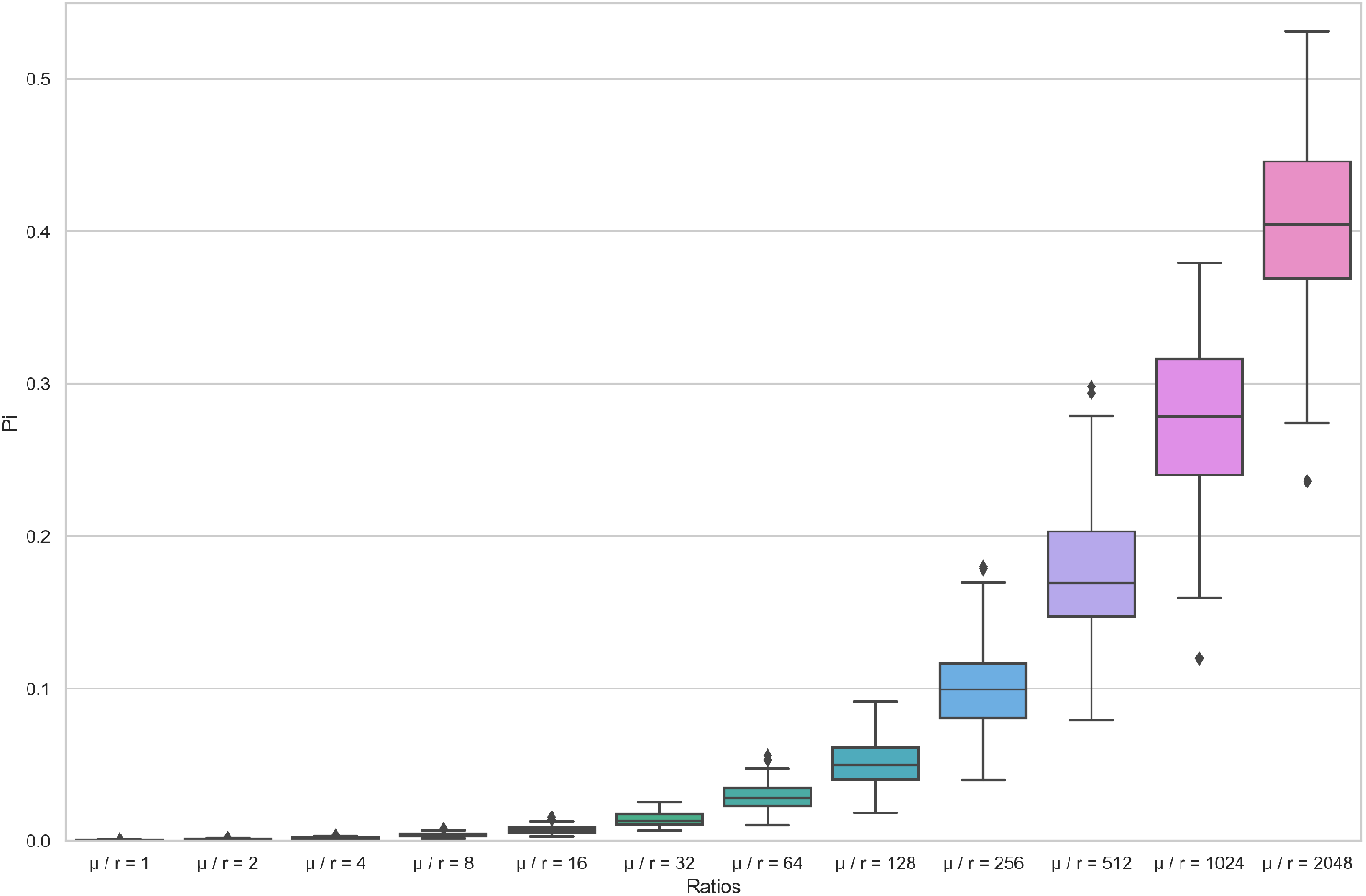
Pi which reflects sequence diversity of simulated sequences under different *µ/r* ratios.

**Figure S3.**
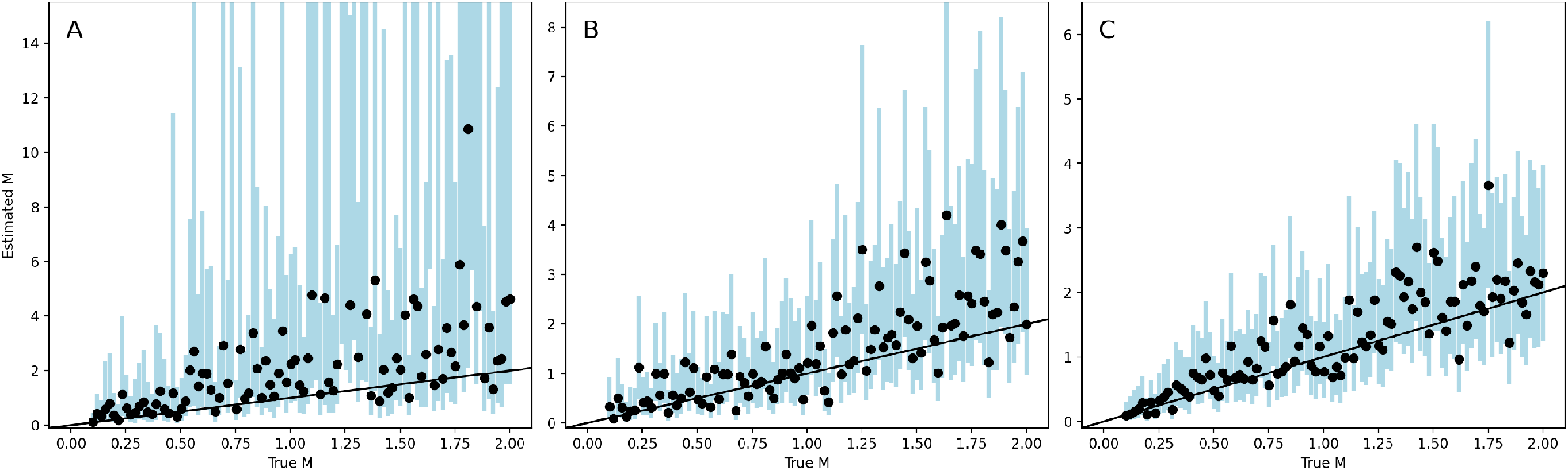
Estimating migration rates M with different sample sizes. Estimating migration rates M between two subpopulations with (A) 20 samples, (B) 50 samples, and (C) 100 samples. Each black line is *x* = *y*. Dots and blue bars represent the median posterior estimates and the 95% confidence intervals for each simulation.

**Figure S4.**
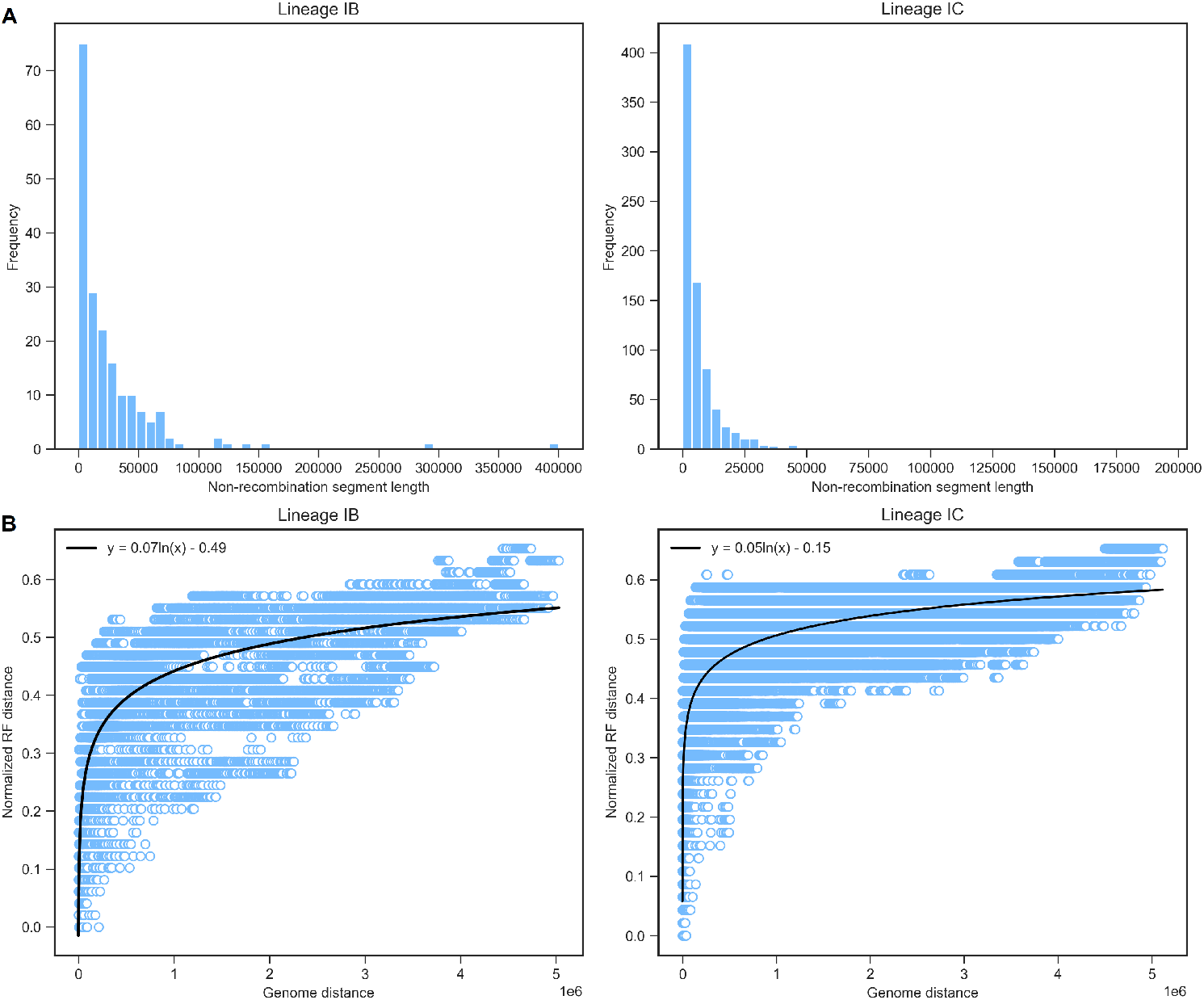
The relationship of genome location distance and RF distance of pairs of local trees, and the histogram of non-recombination segment length (neighboring breakpoints distance). When calculating the genome location distance, we set the middle location of each genome region as coordinates, and then the distance is the absolute value of coordinates difference between two trees.

**Figure S5.**
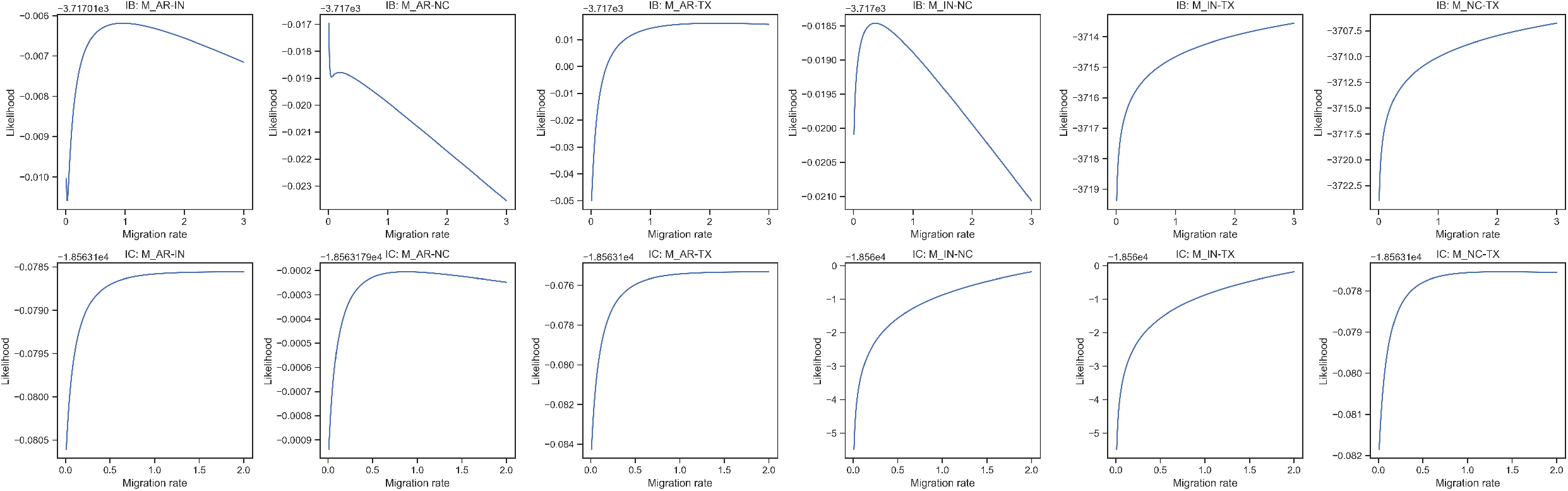
The migration rates (per generation) likelihood profile of lineages IB and IC between subpopulations in four states using the SCAR model.

